# Resveratrol induces mitochondrial dysfunction and decreases chronological life span of *Saccharomyces cerevisiae* in a glucose-dependent manner

**DOI:** 10.1101/118307

**Authors:** Minerva Ramos-Gomez, Ivanna Karina Olivares-Marin, Melina Canizal-García, Juan Carlos González-Hernández, Gerardo M. Nava, Luis Alberto Madrigal-Perez

## Abstract

A broad range of health benefits have been attributed to resveratrol (RSV) supplementation in mammalian systems, including the increases in longevity. Nonetheless, despite the growing number of studies performed with RSV, the molecular mechanism by which it acts still remains unknown. Recently, it has been proposed that inhibition of the oxidative phosphorylation activity is the principal mechanism of RSV action. This mechanism suggests that RSV might induce mitochondrial dysfunction resulting in oxidative damage to cells with a concomitant decrease of cell viability and cellular life span. To prove this hypothesis, the chronological life span (CLS) of *Saccharomyces cerevisiae* was studied as it is accepted as an important model of oxidative damage and aging. In addition, oxygen consumption, mitochondrial membrane potential, and hydrogen peroxide (H_2_O_2_) release were measured in order to determine the extent of mitochondrial dysfunction. The results demonstrated that the supplementation of *S. cerevisiae* cultures with 100 μM RSV decreased CLS in a glucose-dependent manner. At high-level glucose, RSV supplementation increased oxygen consumption during the exponential phase yeast cultures, but inhibited it in chronologically aged yeast cultures. However, at low-level glucose, oxygen consumption was inhibited in yeast cultures in the exponential phase as well as in chronologically aged cultures. Furthermore, RSV supplementation promoted the polarization of the mitochondrial membrane in both cultures. Finally, RSV decreased the release of H_2_O_2_ with high-level glucose and increased it at low-level glucose. Altogether, this data supports the hypothesis that RSV supplementation decreases CLS as a result of mitochondrial dysfunction and this phenotype occurs in a glucose-dependent manner.

## Introduction

Resveratrol (3, 4`, 5-trihydroxystilbene, RSV) is a polyphenol synthetized by plants that has been recognized for its benefits to human health (Jeandet et al. 2014). However, despite the large number of studies identifying phenotypes regulated by RSV, the molecular mechanism by which it acts still remains unknown. There is growing evidence supporting the idea that inhibition of oxidative phosphorylation (OXPHOS) is the mechanism of RSV action (Hawley et al. 2010; Madrigal-Perez and Ramos-Gomez 2016). Several studies indicate that RSV inhibits OXPHOS activity at two sites: in complex In of the electron transport chain (Zini et al. 1998) and the F_1_F_0_-ATPase (Gledhill et al. 2007; Zheng and Ramirez 2000). The inhibition of complex III and the F_1_F_0_-ATPase by RSV could promote a reduced coenzyme Q pool, that enhances reduction of oxygen by complex I, generating a superoxide anion by reverse electron transport (Murphy 2009). In accordance with this hypothesis, supplementation of mammalian cells with RSV resulted in the increased generation of reactive oxygen species (ROS) (Hussain et al. 2011; Plauth et al. 2016; Yoshino et al. 2012). This idea suggests that RSV could induce mitochondrial dysfunction and trigger oxidative damage to cells with a concomitant decrease in cell viability. In mammalian models, reduction of cell viability is a common phenotype provoked by RSV and has been associated with an induction of intrinsic mitochondria-mediated apoptosis (Whitlock and Baek 2012). This can be activated by exacerbated oxidative stress generated by RSV or by the inhibition of F_1_F_0_-ATPase, which reduces mitochondrial ATP production causing decreased sarco/endoplasmic reticulum calcium transport ATPase (SERCA) activity and a mitochondrial calcium overload; that, in turn, initiate the apoptotic pathway (Madreiter-Sokolowski et al. 2016).

In addition, the increase of ROS generation by RSV could have a negative impact on the aging process as it has been postulated in the free radical theory of aging (Harman 1956). In accordance with this hypothesis, aging studies showed that RSV fails to extend life span in *Saccharomyces cerevisiae* (Howitz et al. 2003; Kaeberlein et al. 2005), mosquito *Anopheles stephensis* (Johnson and Riehle 2015) and *Drosophila melanogaster* (Bass et al. 2007). Epidemiology evidence even suggests that RSV effect is dependent of diet (Yoshino et al. 2012). In this regard, recent studies have started to highlight that cellular changes induced by RSV could be influenced by the cellular energy status (Cai et al. 2015; Madrigal-Perez et al. 2016).

Thus, the aim of this study was to prove that RSV decreases cellular life span by mitochondrial dysfunction in a glucose-dependent manner (main source of carbon and energy in the growth media). *S. cerevisiae* was used because it is recognized as an important model for exploring mitochondrial processes, aging, and RSV mechanisms (Kaeberlein et al. 2005; Madrigal-Perez et al. 2015). Moreover, chronological life span (CLS) of *S. cerevisiae* has been accepted as an key model of oxidative damage and aging (Kaeberlein 2010). In this study, it was found that RSV provoked the decrease of CLS in *S. cerevisiae* in a glucose dependent-manner, and this effect is related with the mitochondrial dysfunction instigated by RSV.

## Material and Methods

### Strains

The experiments were performed using *S. cerevisiae* BY4742 (Matalpha; *his3Δ1*; *leu2Δ0*; *lys2Δ0*; *ura3Δ0*) acquired from EUROSCARF (Frankfurt, Germany). The strain was maintained in yeast extract-peptone-dextrose (YPD) medium (1% yeast extract, 2% casein peptone and 2% glucose).

### Chronological life span

The CLS was determined according to Murakami and Kaeberlein (2009) with some modifications. Synthetic-complete (SC) medium was prepared and consisted of 0.18% yeast nitrogen base without amino acids (Sigma-Aldrich, St. Louis, MO, USA), 0.5% ammonium sulfate (JT Baker, Center Valley, PA, USA), 0.2% KH_2_PO_4_ (JT Baker), 1% drop-out mix without uracil (Sigma-Aldrich), supplemented with 400 μg/mL of uracil (Sigma-Aldrich). The mediums had three different levels of glucose (0.5, 2 and 10% glucose; JT Baker) and five levels of RSV (0, 10, 30, 50 and 100 μM; Sigma-Aldrich) that were added at the beginning of the CLS assay. Posteriorly, 2 mL of the SC medium were inoculated with 1% of a fresh overnight culture of *S. cerevisiae* in a test tube with 20% headspace volume. The cultures were grown at 30 °C with shaking (180 rpm) for 15 days. After three days of incubation, aliquots of 5 μL were taken and inoculated into 145 μL YPD 2% glucose, every two days. Samples were placed in a honeycomb plate (Growth Curves, Piscataway, NJ, USA) and incubated at 30 °C for 24h in a Bioscreen (model C MBR, Growth Curves) programmed with continuous shaking and readings at 600 nm each 30 min. Finally, the survival percentage (*Sn*) was calculated according to equation 1:

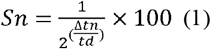

Where *Δtn* is the time shift (h), obtained by interpolation of D.O._600_=0.5 using a linear regression of the exponential phase of growth curve, and *td* is the doubling time (h).

### Determination of *in situ* mitochondrial respiration

The oxygen consumption was measured polarographically at 28 °C using a Clark detector (YSI, model 5,300, Yellow Springs, OH, USA). The *S. cerevisia* cultures were harvested at mid-log phase (D.O. _600_ ~ 0.5; exponential phase yeast culture) or fifteen days after inoculation (chronologically aged yeast culture) at 5000 × *g* for 5 minutes at 28 °C. Subsequently, 125 mg of cells were resuspended in 5 mL of buffer 10 mM 2-(N-morpholino) ethanesulfonic acid (MES-TEA; Sigma-Aldrich), adjusted to pH 6.0 with triethanolamine (Sigma-Aldrich) and placed in a closed chamber with constant stirring. The basal respiration was determined using 10 mM glucose as a substrate. The maximal rate of mitochondrial respiratory chain function was evaluated with the addition of the mitochondrial uncoupler CCCP (10 μM; Sigma-Aldrich). The oxygen consumption due to non-mitochondrial sources was determined using 10 μM Antimycin A (Sigma-Aldrich) and was subtracted to the basal respiration and maximal respiratory capacity values. The spare respiratory capacity (SRC) was calculated from the maximal respiratory capacity minus basal respiration. Oxygen consumption was expressed as nanoatoms/(min)(mg of cells).

### Measurement of mitochondrial membrane potential *in situ*

An important mediator of OXPHOS activity is the mitochondrial membrane potential. Therefore, the mitochondrial membrane potential measurement can give a better understanding of the effect of RSV on OXPHOS activity. For the determination of mitochondrial membrane potential, *S. cerevisia* cultures at mid-log phase (D.O. _600_ ~ 0.5; exponential phase yeast culture) or fifteen days after inoculation (chronologically aged yeast culture) that were harvested at 5000 × *g* for 5 minutes at 28 °C were used. The cells were resuspended in 2 mL of citric acid-Na_2_HPO_4_ buffer adjusted to pH 7 containing 100 nM DiSC_3_, 20 mM glucose and 200 μM CaCl_2_. Subsequently, cells were plated in 96- well black plate at a density of 3 × 10^6^ cells/well. The fluorescence signal was measured at an excitation wavelength of 572 nm and an emission of 582 nm with a microplate reader (Varioskan flash spectral scanning multimode reader, Thermo-Scientific). First, the background signal for 50 seconds was measured. Posteriorly, CCCP stock solution was added to obtain 10 μM as a final concentration in each well and incubated in the dark at 28 °C for 30 minutes with constant stirring.

### Quantification of H_2_O_2_ release

Hydrogen peroxide (H_2_O_2_) release was used as an indicator of ROS generation and was determined by the Amplex red hydrogen peroxide assay kit (Invitrogen, Waltham, MA, USA) according to manufacturer instructions. Overnight *S. cerevisia* cultures and cultures that had been inoculating for fifteen days (chronologically aged yeast cultures) were harvested at 5000 × *g* for 5 minutes at 28 °C. The cells were resuspended in 2 mL of assay buffer containing 20 mM Tris-HCl, 0.5 mM EDTA, 2% of ethanol at pH 7. The cells were plated into a black plate at a density of 3 × 10^6^ cells/well. The basal release of H_2_O_2_ was measured at an excitation wavelength of 563 nm and an emission of 587 nm with a microplate reader (Varioskan, Thermo-Scientific).

### Statistical analyses

To analyze results of CLS under different levels of RSV and glucose, repeated one-way ANOVA measures followed by Dunnett’s test were used. One-way ANOVA was applied to test for the differences in oxygen consumption rate, mitochondrial membrane potential, and H_2_O_2_ release, followed by Dunnett’s test. The rest of the experiments were analyzed using ANOVA with Tukey’s test. For all analyses, at least 4 independent experiments with 3 technical repetitions were performed. Statistical analyses were computed using GraphPad Prism 6.00 for Macintosh (GraphPad Software, La Jolla California, USA).

## Results

### Resveratrol decreases chronological life span in a glucose-dependent manner

First, the effect of RSV supplementation on CLS and its relationship with glucose availability was established. For this purpose, the percentage of cell viability of *S. cerevisia* cultures supplemented with five levels of RSV (0, 10, 30, 50 and 100 μM) and three levels of glucose (0.5, 2 and 10% glucose), for 12 days was measured. Therefore, under this model, 0.5% of glucose corresponds to low-level (caloric restriction), 2% to standard-level and 10% to high-level of glucose. It is important to mention that the growth of *S. cerevisiae* was only slightly higher in cultures grown with 2% glucose (specific growth rate, μ~0.2379 h^−1^) in comparison to cultures grown with 10% glucose (μ~0.2019 h^−1^), whereas at 0.5% glucose, *S. cerevisiae* displayed the slower growth (μ~0.1602 h^−^) (*P*>0.0001; Fig. 1). This data corroborates the physiological differences among the three concentrations of glucose used in this study.

**Fig. 1.**
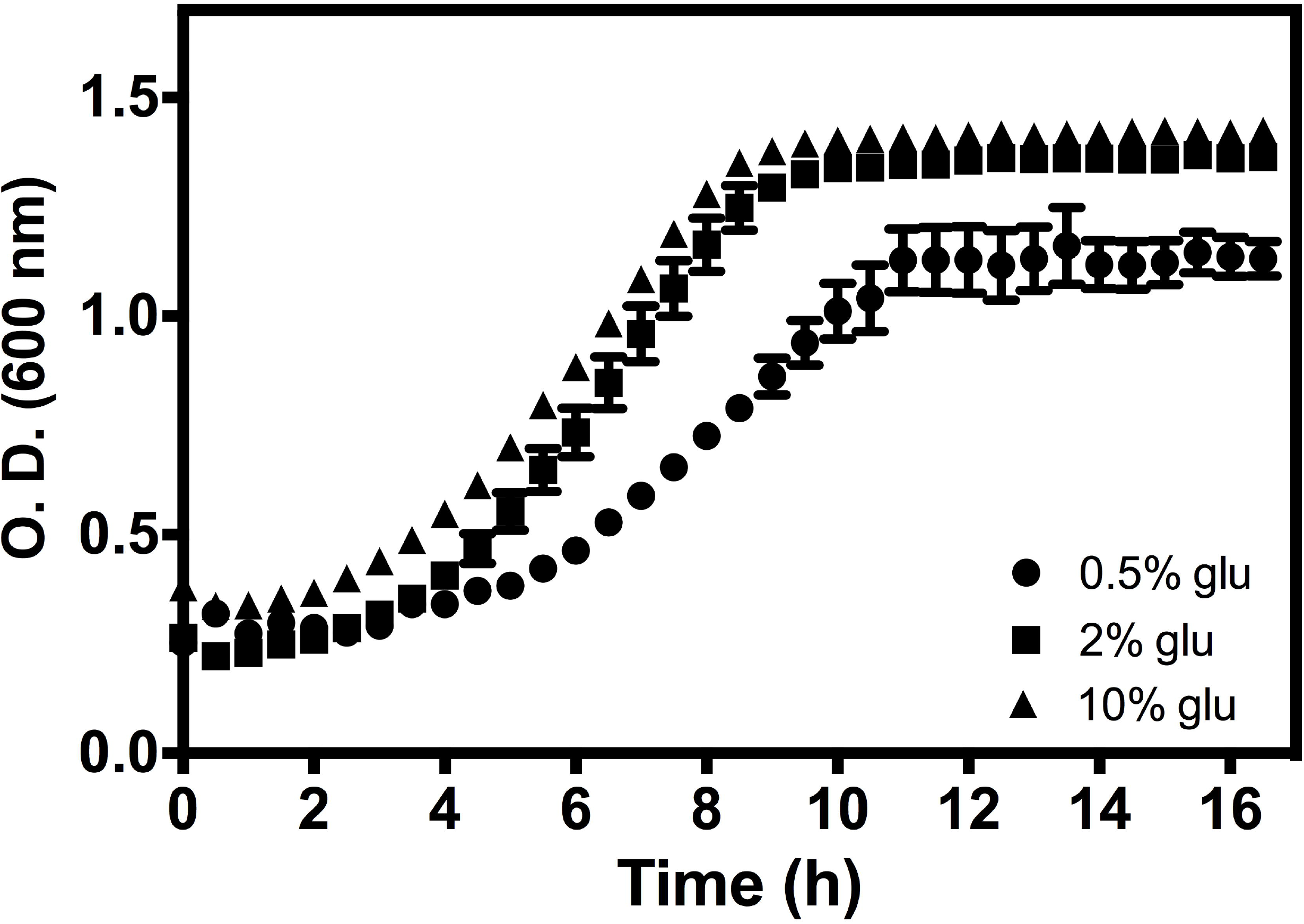
Effect of glucose concentration on *S. cerevisia* growth. The cell growth curve of *S. cerevisia*growth with 0.5, 2 and 10% of glucose. The results represent mean values ± SEM from 4 independent experiments, which includes mean values of 3 technical repetitions. Statistical significance was calculated by repeated measures one-way ANOVA followed by Tukey’s test (*P*<0.0001).

By taking advantage of this model, it would found that cultures of *S. cerevisiae* supplemented with the high-level glucose and RSV showed no significant differences in CLS with respect to control (without RSV) (*P*>0.05; Fig. 2a). We did not observe any differences in CLS between *S. cerevisia* cultures grown with 2% glucose (standard-level) without RSV (control) and those supplemented with 10, 30, and 50 μM RSV. However, the cultures supplemented with 100 μM RSV had a decrease in CLS, when compared to the control (*P*<0.05; Fig. 2b). At the low-level glucose, only cultures supplemented with 100 μM RSV showed a decrease in CLS, in comparison to the control (*P*<0.05; Fig. 2c). Altogether, this data showed that RSV supplementation fails to extend CLS of *S. cerevisia* A negative effect in CLS was evident in cultures supplemented with 100 μM RSV. Indeed, the decrease of cell viability by 100 μM RSV is glucose dose-dependent (*P*<0.05; Fig. 3). Overall, this data supports the hypothesis that RSV could induce a cellular oxidative damage with a concomitant decrease in life span and cellular viability. Importantly, this last phenotype is only evident at standard and lower concentrations of glucose.

**Fig. 2.**
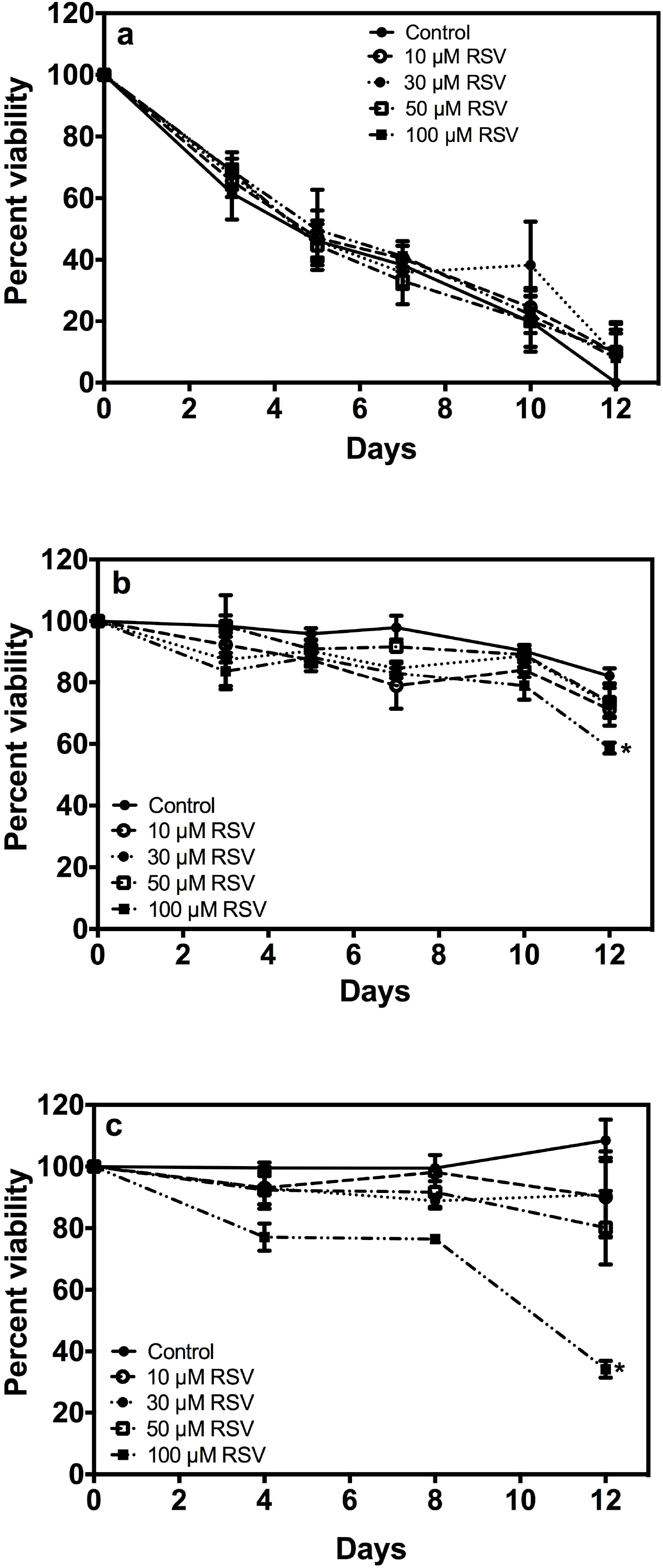
Influence of resveratrol in chronological life span (CLS) of *S. cerevisiae* at different glucose concentrations. The CLS of *S. cerevisiae* was assayed in minimal medium (SC) supplemented with: **a)** 10% glucose; **b)** 2% glucose; **c)** 0.5% glucose. Results represent mean values ± SEM from 4 independent experiments, which includes mean values of 3 technical repetitions. Statistical significance was calculated by repeated measures one-way ANOVA followed by Dunnett's test (*\*P*<0.05 vs. control).

**Fig. 3.**
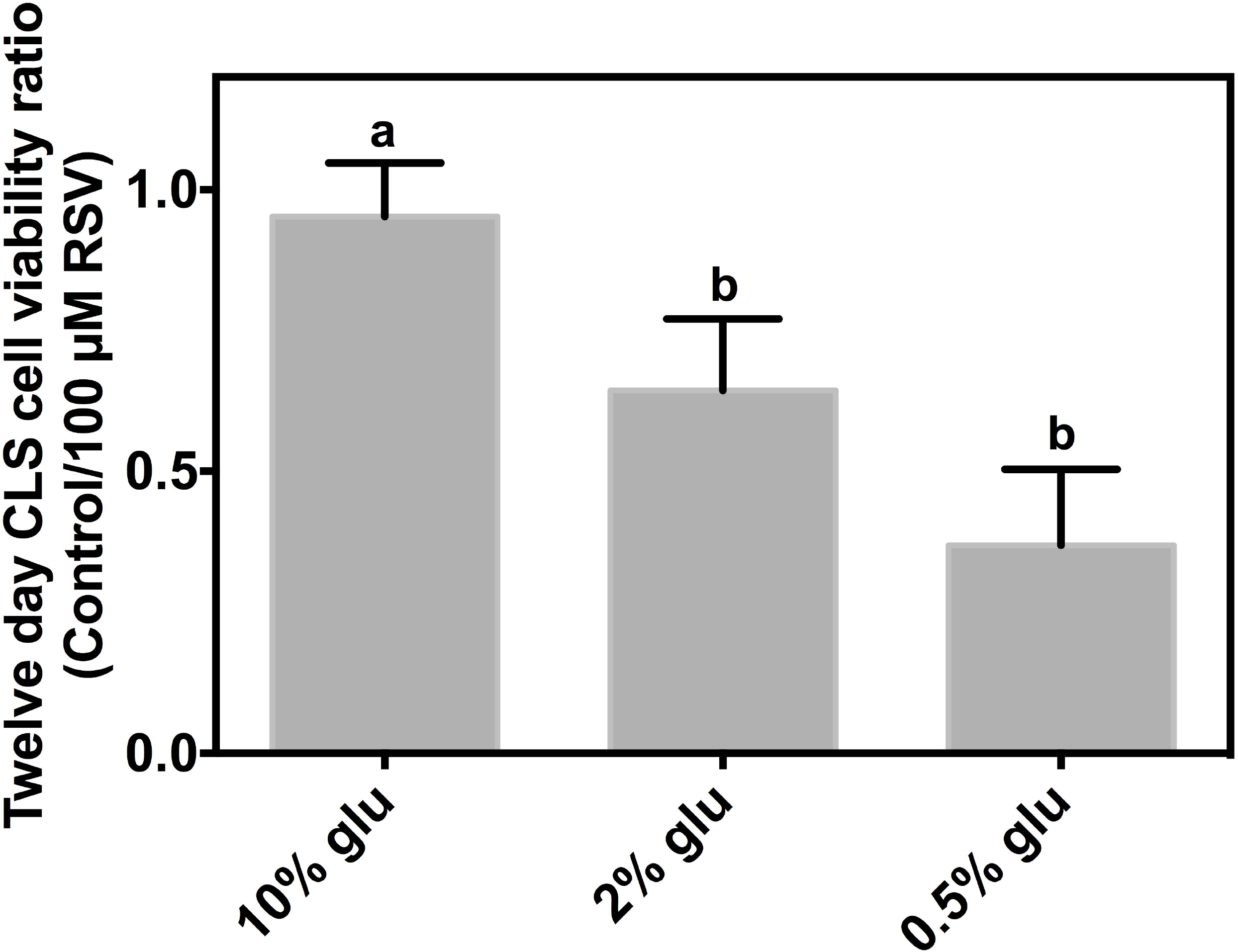
Resveratrol effect on *S. cerevisiae* cellular viability is glucose dependent. The cellular viability of *S. cerevisia* cultures supplemented with 10, 2 and 0.5% glucose (glu) and 100 μM resveratrol (RSV) at 12 days of chronological aging was calculated by dividing the doubling time of RSV supplemented cells between the doubling time of control cells (without RSV). Results represent mean values ± SEM obtained from 4-8 independent experiments with three technical replicates each. Statistical analyses were performed using one-way ANOVA followed by Tukey’s test (*P*≤0.05). Different letters indicates statistical differences.

### Effect of resveratrol on mitochondrial respiration is glucose dose-dependent

Oxygen consumption is an indicator of the function of the electron transport chain and proton leakage caused by the F_1_F_0_-ATPase. Therefore, in order to evaluate whether the decrease of cell viability by RSV at standard and lower concentrations of glucose is related to OXPHOS activity, oxygen consumption analysis was performed in yeast cultures in the exponential phase and chronologically aged (15 days of age) with the same levels of RSV and glucose used in the CLS assay.

Unexpectedly, exponential phase yeast cultures of *S. cerevisiae* grown in high-level glucose and supplemented with RSV showed an increase of basal respiration and the maximal respiratory capacity in a dose-dependent manner (*P*<0.0001; Fig. 4). At the standard-level glucose, no differences were detected with RSV supplementation, except at high-level of RSV (100 μM), which prompts basal respiration and the maximal respiratory capacity (*P*<0.05; Fig. 4). Interestingly, RSV supplementation inhibited basal respiration and the maximal respiratory capacity of *S. cerevisia* cultures grown at low-level glucose (*P*<0.0001; Fig. 4). With respect to spare respiratory capacity (SRC), it was observed that *S. cerevisia* cultures grown at high-level glucose increased the SRC only when they were supplemented with 10 μM RSV (*P*<0.0001; Fig. 4). However, SRC was decreased with all RSV levels assayed at low-level glucose (*P*<0.0001; Fig. 4). Remarkably, at a low-level glucose it was evident that there was a drastic inhibition of basal respiration as well as in the maximal respiratory capacity. This was accompanied by a decrease in the SRC with RSV supplementation, which indicates a relationship between inhibition of mitochondrial respiration and cell viability in early stages of growth.

**Fig. 4.**
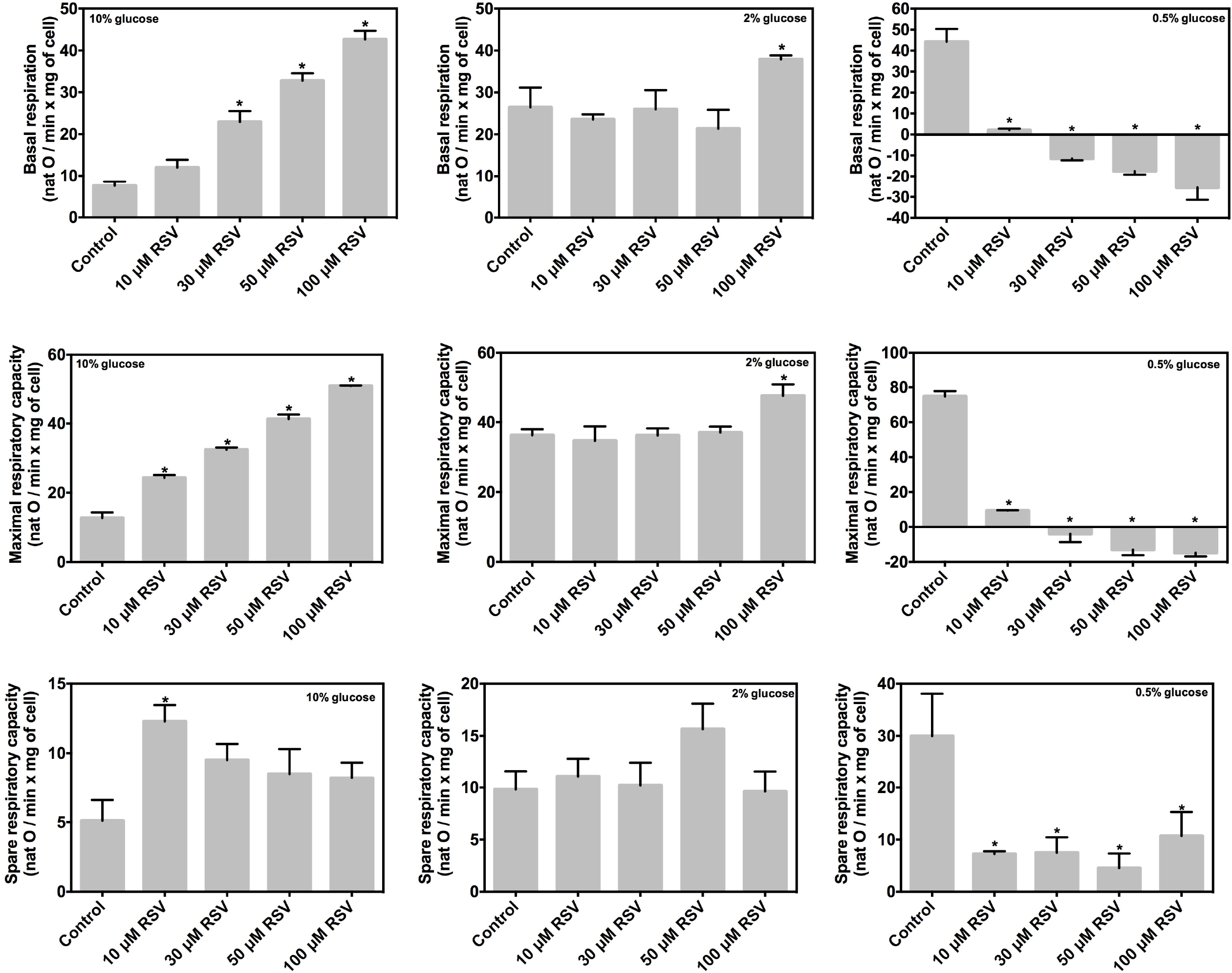
The influence of resveratrol (RSV) on oxygen consumption of exponential phase *S. cerevisia*cultures is glucose dependent. The oxygen consumption was measured at basal state and maximal respiratory capacity in *S. cerevisia* cells grown in YPD medium with 10, 2 and 0.5% glucose and supplemented with 10, 30, 50 and 100 μM RSV. The spare respiratory capacity (SRC) was calculated from the maximal respiratory capacity minus basal respiration. Results represent mean values ± SEM from 3 independent experiments. Statistical analyses were performed using one-way ANOVA followed by Dunnett’s test (*\*P*≤0.05 vs. control).

On the other hand, chronologically aged yeast cultures at high-levels of glucose and supplemented with RSV showed an inhibition of basal respiration and the maximal respiratory capacity (*P*<0.0001; Fig. 5). Interestingly, the control culture (without RSV) grown at high-level glucose increased basal respiration and the maximal respiratory capacity in comparison to exponential phase yeast cultures grown under the same conditions. This is probably due to the post-diauxic phase, where *S. cerevisia*maintains cell functions using mitochondrial respiration, by oxidizing by-products of fermentation (e.g. ethanol and acetic acid). At standard-level of glucose, *S. cerevisiae* did not display any difference in basal respiration or the maximal respiratory capacity (*p*<0.0001; Fig. 5). Finally, in *S. cerevisia*cultures grown in low glucose conditions, an inhibition of basal respiration as well as maximal respiratory capacity was observed (*p*<0.0001; Fig. 5). These results confirm the previous observation that RSV supplementation inhibits mitochondrial respiration in post-diauxic phase (Madrigal-Perez et al. 2016), which supports the idea that OXPHOS is the main target of RSV.

**Fig. 5.**
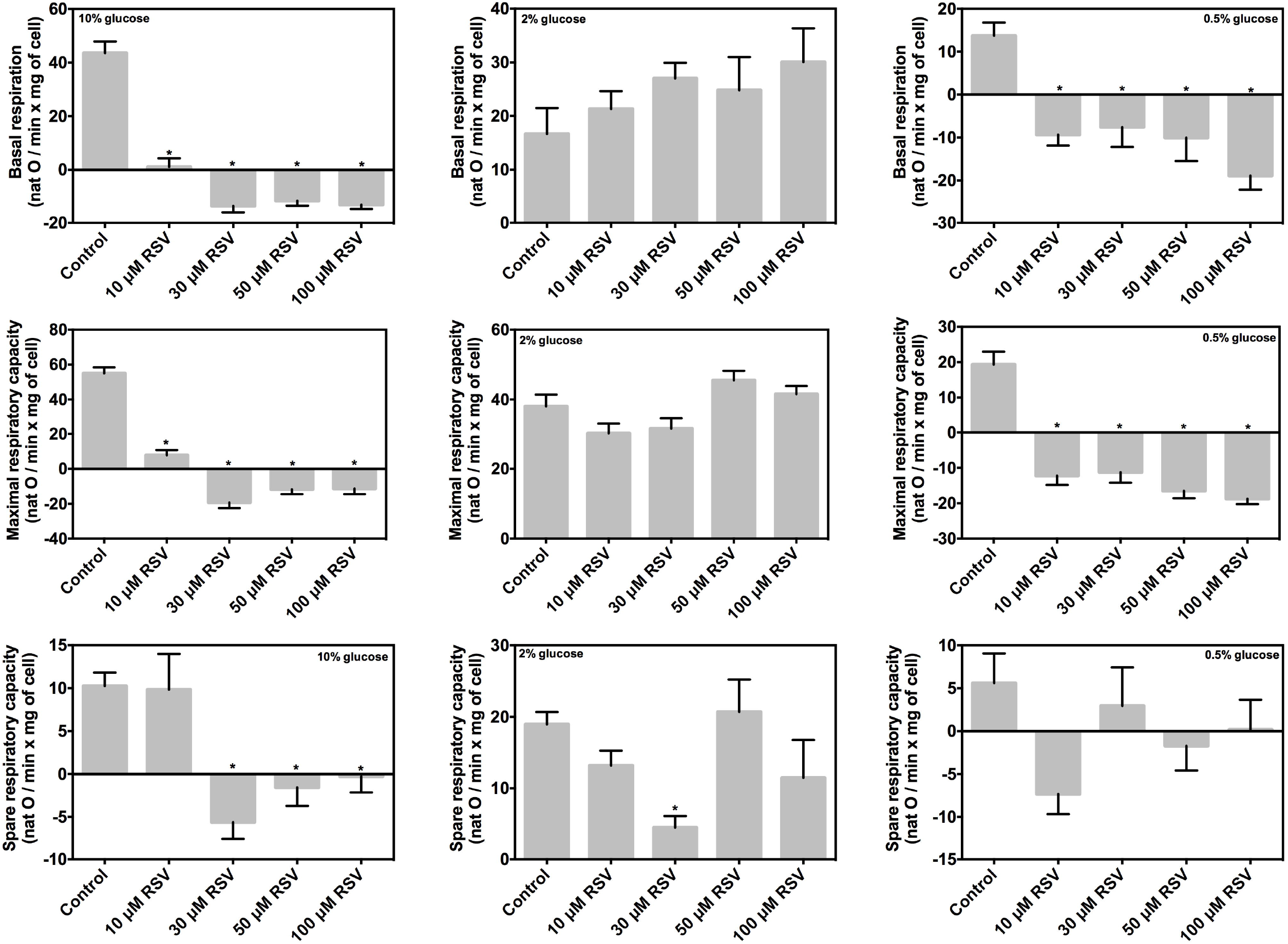
The influence of resveratrol (RSV) on oxygen consumption of chronologically aged *S. cerevisia* cultures is glucose dependent. The oxygen consumption was measured at basal state and maximal respiratory capacity in *S. cerevisia* cells grown in YPD medium with 10, 2 and 0.5% glucose and supplemented with 10, 30, 50 and 100 μM RSV. The spare respiratory capacity (SRC) was calculated from the maximal respiratory capacity minus basal respiration. Results represent mean values ± SEM from 3 independent experiments. Statistical analyses were performed using one-way ANOVA followed by Dunnett test (*\*P*≤0.05 vs. control).

Overall, these results indicate that RSV supplementation impairs oxygen consumption in low-glucose conditions in exponential phase yeast cultures, which probably exerts a negative impact in cell viability and CLS. In addition, RSV supplementation inhibits oxygen consumption of *S. cerevisia* cultures grown in high-level glucose but only when those enter in the post-diauxic phase. However, exponential phase yeast cultures grown in high-level glucose and supplemented with RSV increased oxygen consumption, this probably due to a mitochondrial membrane impairment.

### Resveratrol supplementation disturbs the mitochondrial membrane potential

The mitochondrial membrane potential was measured in order to corroborate whether the effect on oxygen consumption occasioned by RSV is due to an alteration of mitochondrial membrane potential.

In exponential phase yeast cultures, it was found that supplementation with 30 and 50 μM RSV in cultures grown at high-level glucose increased mitochondrial membrane potential in comparison to the control (*P*<0.05; Fig. 6). In the case of the *S. cerevisia* cultures grown with 2% glucose (standard-level), the supplementation with 50 μM RSV showed an increase in the mitochondrial membrane potential (*P*<0.05; Fig. 6). Finally, at low-level glucose only a 100 μM of RSV increased the mitochondrial membrane potential (*P*<0.05; Fig. 6). On the contrary, in chronologically aged yeast cultures (15 days of age), it was observed that *S. cerevisia* cultures grown at low-level glucose increased the mitochondrial membrane potential when they were supplemented with 10 and 30 μM RSV (*P*<0.05; Fig. 7). Nonetheless, no differences in mitochondrial membrane potential were observed in cells grown in standard and high-levels of glucose and supplemented with RSV compared to controls (*P*>0.05; Fig. 7).

**Fig. 6.**
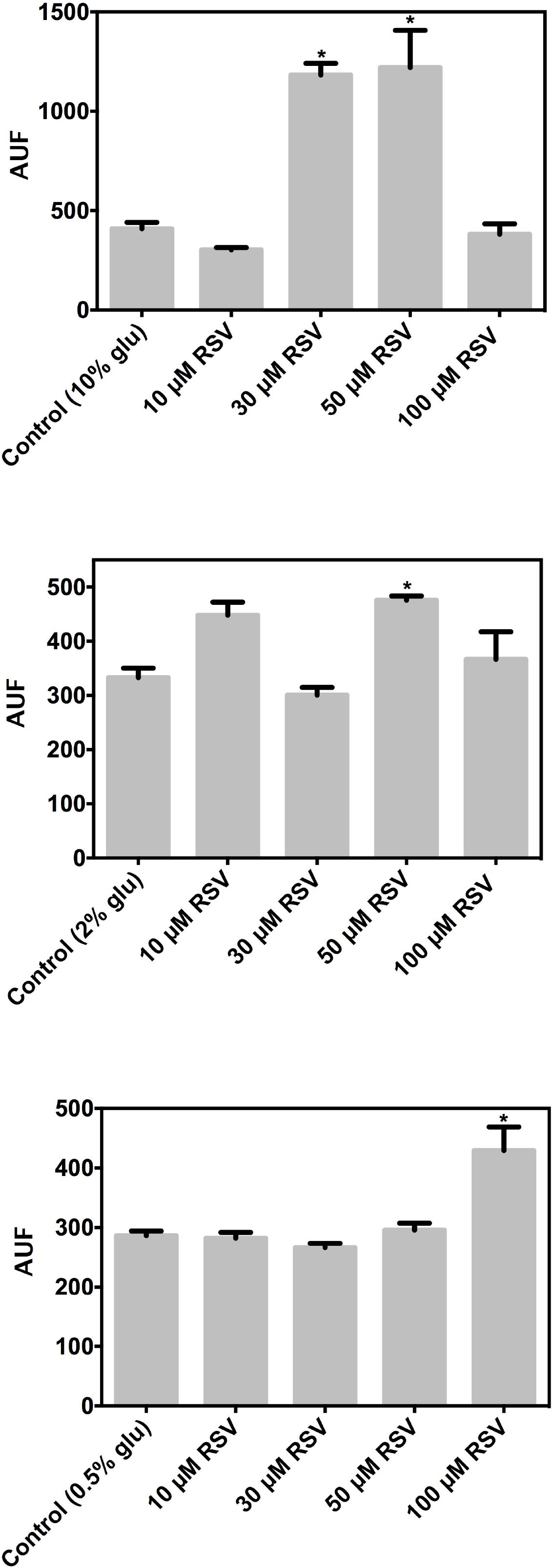
The influence of resveratrol (RSV) on mitochondrial membrane potential in exponential phase *S. cerevisia* cells. The mitochondrial membrane potential was measured by DiSC_3_ quenching assay in *S. cerevisia* cells grown in YPD medium with 10, 2, and 0.5% glucose and supplemented with 10, 30, 50 and 100 μM RSV. Results represent mean values ± SEM from 5 independent experiments. Statistical analyses were performed using one-way ANOVA followed by Dunnett’s test (*\*P*≤0.05 vs. Control cells).

**Fig. 7.**
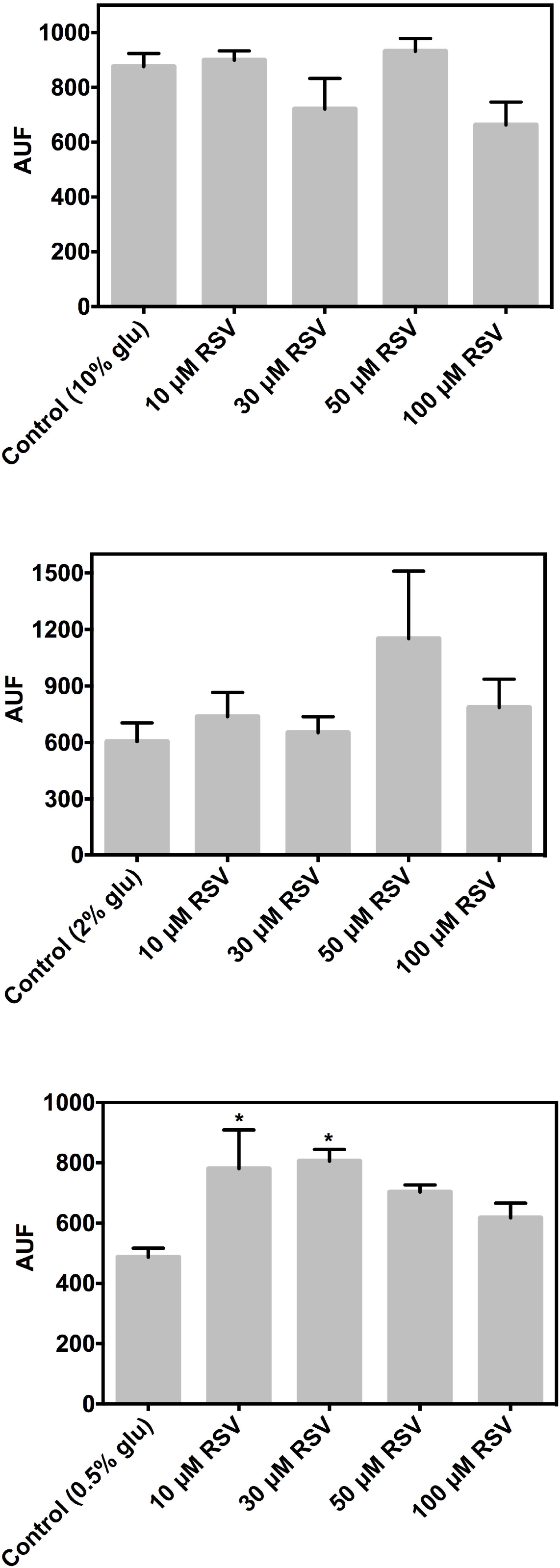
The influence of resveratrol (RSV) on mitochondrial membrane potential in chronologically aged *S. cerevisia* cells. The mitochondrial membrane potential was measured by DiSC_3_ quenching assay in *S. cerevisia* cells grown in YPD medium with 10, 2, and 0.5% glucose and supplemented with 10, 30, 50 and 100 μM RSV. Results represent mean values ± SEM from 5 independent experiments. Statistical analyses were performed using one-way ANOVA followed by Dunnett’s test (*\*P*≤0.05 vs. Control cells).

Altogether, this data suggests that RSV could act as inhibitor of the proton leakage, probably by inhibiting the F_1_F_0_-ATPase activity as it has been previously proposed.

### Influence of resveratrol in the release of H_2_O_2_

The effect of RSV on oxygen consumption and mitochondrial membrane potential suggests that ROS generation could be disturbed by RSV supplementation. For this reason, the release of H_2_O_2_ as an indicator of the ROS generation was measured.

Exponential phase yeast cultures grown at high-level glucose decreased the release of H_2_O_2_ in all the concentrations of RSV tested with respect to control (*P*<0.05; Fig. 8). At standard-level glucose only the highest level of RSV (100 μM) decreased release of H_2_O_2_ (*P*<0.05; Fig. 8). Interestingly, RSV supplementation at 10 μM increased the release of H_2_O_2_ at low-level glucose, whereas no significant effect was observed at higher RSV levels (*P*<0.05; Fig. 8). On the contrary, chronologically aged yeast cultures grown at low-level glucose decreased the release of H_2_O_2_ when they were supplemented with 10, 50 and 100 μM RSV (*P*<0.05; Fig. 9); whereas no differences in the release of H_2_O_2_ of *S. cerevisiae* cultures grown in standard and high concentrations of glucose and supplemented with RSV at any level were observed (*P*>0.05; Fig. 9). Altogether, this data corroborates the bioenergetics effect of RSV on OXPHOS.

**Fig. 8.**
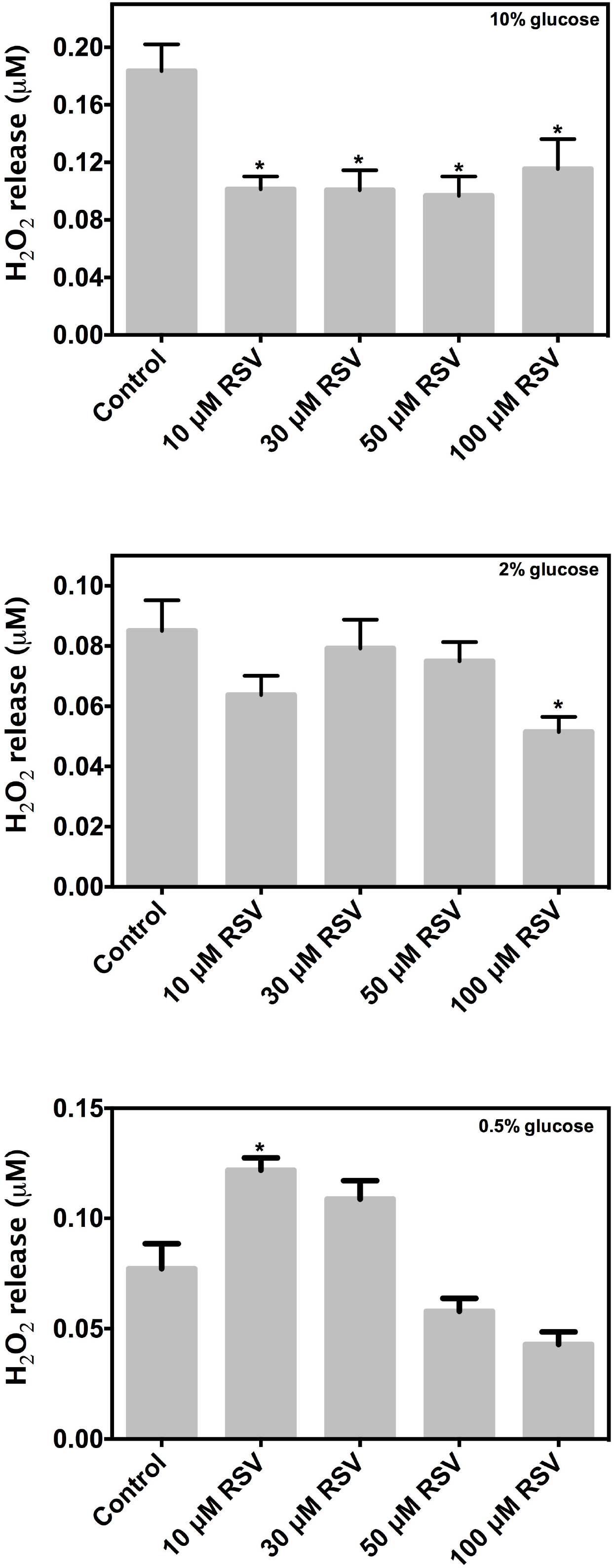
The influence of resveratrol (RSV) on H_2_O_2_ release in exponential phase *S. cerevisia* cells. The H_2_O_2_ release was measured by Amplex red kit in *S. cerevisia* cells grown in YPD medium with 10, 2, and 0.5% glucose and supplemented with 10, 30, 50 and 100 μM RSV. Results represent mean values ± SEM from 5 independent experiments. Statistical analyses were performed using one-way ANOVA followed by Dunnett’s test (*\*P*≤0.05 vs. Control cells).

**Fig. 9.**
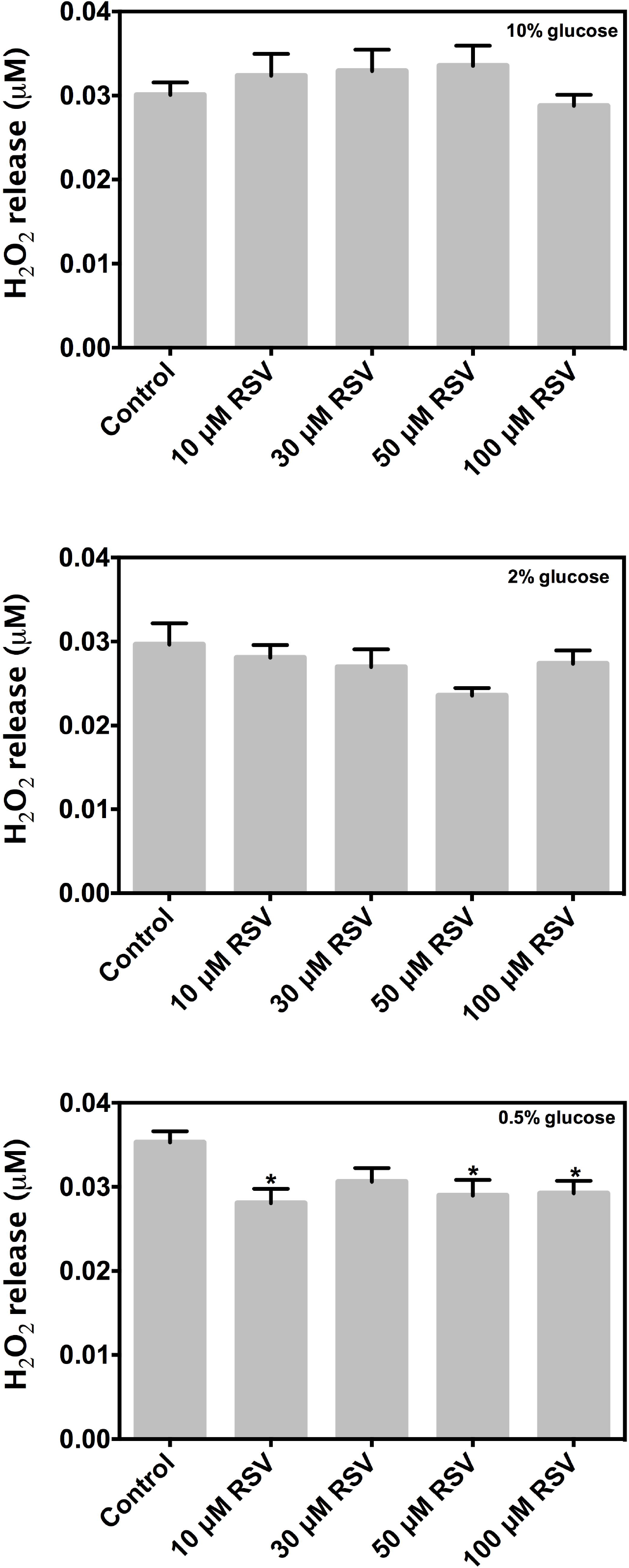
The influence of resveratrol (RSV) on H_2_O_2_ release in chronologically aged *S. cerevisia* cells. The H_2_O_2_ release was measured by Amplex red kit in *S. cerevisia* cells grown in YPD medium with 10, 2, and 0.5% glucose and supplemented with 10, 30, 50 and 100 μM RSV. Results represent mean values ± SEM from 5 independent experiments. Statistical analyses were performed using one-way ANOVA followed by Dunnett’s test (*\*P*≤0.05 vs. Control cells).

## Discussion

Despite the large amount of RSV studies in the biomedical field, the molecular mechanism by which it acts is uncertain. It has recently been proposed that inhibition of the OXPHOS activity is the mechanism of RSV action (Hawley et al. 2010; Madrigal-Perez and Ramos-Gomez 2016). However, additional evidence is required to solidify this idea. Herein, we provide evidence that RSV reduces CLS by induction of mitochondrial dysfunction. First, the study demonstrates that RSV fails to extend CLS and high-levels of RSV (100 μM) decrease CLS in a glucose-dependent manner. Second, at high-level glucose RSV supplementation increases oxygen consumption in exponential phase yeast cultures, but inhibits it in chronologically aged yeast cultures. However, at low-level glucose, oxygen consumption was inhibited in both yeast cultures, exponential phase and chronologically aged. Third, RSV supplementation promotes the polarization of the mitochondrial membrane. Fourth, in the exponential phase yeast cultures, the H_2_O_2_ release decreases by RSV supplementation at high-level glucose, whereas, at low-level glucose H_2_O_2_ release increases. However, in chronologically aged yeast cultures, the H_2_O_2_ release decreases by RSV supplementation in low-level glucose and remains unchanged in high-level glucose. Collectively, this data supports the hypothesis that the main target of RSV is the OXPHOS, and suggests that RSV shortens CLS by this mechanism in a glucose-dependent manner.

There is conflicting data concerning the extension of life span in *S. cerevisiae* by RSV supplementation. Early studies demonstrated that RSV supplementation (10, 100 and 500 μM) extends replicative life span in *S. cerevisiae* (Howitz et al. 2003). However, other studies have shown that RSV supplementation at similar doses (10 and 100 μM) does not extend replicative life span of the strains BY4742 and W303 (Kaeberlein et al. 2005). On the other hand, the CLS of *S. cerevisiae* was decreased by RSV (2 mg/L) supplementation in winemaking conditions (synthetic grape juice MS300) (Orozco et al. 2012). Nonetheless, *S. cerevisia* cultures grown in standard media (YPD supplemented with 2% glucose) and supplemented with RSV (logarithmic doses ranged from 1 nM to 1 mM) did not show any effect in the CLS (Choi et al. 2013). Interestingly, it was found that in minimum media, RSV supplementation (100 μM) shortens CLS in standard and low-level glucose but not in high-level glucose. A diet-dependent effect on life span by RSV supplementation has also been reported in other organisms. For example, a decrease in survivorship in blood-fed mosquitoes fed with RSV (50, 100 and 200 μM) was found (Johnson and Riehle 2015). On the contrary, RSV supplementation only extends the life span of mice fed with a high-fat diet (Baur et al. 2006) and not with a standard-diet (Pearson et al. 2008). Altogether, this data indicates that RSV effects on life span are dependent on diet composition, suggesting a bioenergetic origin of RSV-induced mechanism.

On the other hand, typical markers of programmed cell death accompanied the decrease in CLS caused by RSV. For example, the phenotype exerted by pheromone and amiodarone, which are well-known inducers of programed death in *S. cerevisiae*, is similar to that found with RSV supplementation in this study. Both experiments were characterized by an increase of mitochondrial respiration, mitochondrial membrane potential, and ROS generation in early stages (exponential cultures in the case of RSV supplementation) (Pozniakovsky et al. 2005). Importantly, supplementation with high levels of amiodarone (40-80 μM) increased mitochondrial respiration. Whereas, a low level of amiodarone (5 μM) decreased mitochondrial respiration as with RSV (Pozniakovsky et al. 2005). In addition, it was found that amiodarone increases the mitochondrial membrane potential, which in turn, increases ROS generation. Importantly, the increase in mitochondrial respiration and mitochondrial membrane potential elicited by amiodarone triggered the rise of the cytoplasmic calcium (Pozniakovsky et al. 2005). Recently, it has been demonstrated that inhibition of the F_1_F_0_-ATPase by RSV reduces mitochondrial ATP production provoking decreased sarco/endoplasmic reticulum calcium transport ATPase (SERCA) activity and mitochondrial calcium overload, which initiates the apoptotic pathway (Madreiter-Sokolowski et al. 2016). Overall, this data suggests that RSV decreases CLS by induction of programmed death in *S. cerevisia* In this regard, this phenotype would be elicited by cytoplasmic and mitochondrial calcium overload, however, additional experiments are needed to confirm this.

A growing number of studies have determined that RSV supplementation disrupts the activity of the electron transport chain between complex I (Moreira et al. 2013), and III and in the F_1_F_0_-ATPase (Gledhill et al. 2007). This phenotype exerts a direct impact on oxygen consumption within cells. For example, it has been reported that RSV (50 μM) supplementation decreased oxygen consumption in mouse colon cancer CT-26 cells (Sassi et al. 2014) and in human embryonic kidney cells supplemented with 300 μM RSV (Hawley et al. 2010). Interestingly, low doses of RSV (1-5 μM) increase mitochondrial complex I activity in liver cells (Desquiret-Dumas et al. 2013). In this regard, *in vitroin* studies demonstrated that RSV competes with NAD, which in turn stimulates the activity of complex I at low doses (<5 μM), whereas high doses (50 μM) it decreased (Gueguen et al. 2015). Similarly, we found that RSV supplementation promotes and inhibits oxygen consumption in a glucose-dependent manner. At high-level of glucose RSV increases oxygen consumption, whereas at low-level of glucose RSV inhibits oxygen consumption in exponential phase yeast cultures. Additionally, data from the present study suggests that RSV supplementation could affect cell viability when cells were under mitochondrial respiration to support their growth (e.g. caloric restriction in *S. cerevisiae*) and this phenotype is more evident at early stages of growth (*S. cerevisiae* exponential phase). In addition, a decrease in the SRC in exponential phase yeast cultures grown at low-glucose level was found. This indicates that the yield of ATP that can be achieved by using mitochondrial respiration as a main pathway for energy production in response to a sudden energy demand (Brand and Nicholls 2011) is restricted by RSV supplementation. Importantly, the phenotype exerted by RSV supplementation in mitochondrial respiration is maintained in the long-term, as shown in chronologically aged yeast cultures grown in high-level of glucose, which presented an inhibition of oxygen consumption in the post-diauxic phase (respiratory phase). Overall, this data indicates that inhibition of oxygen consumption is due to inhibition of OXPHOS activity.

The inhibition of the F_1_F_0_-ATPase proton leakage by RSV suggests that RSV supplementation should increase the mitochondrial membrane potential. Accordingly, it was found that RSV supplementation promotes the polarization of the mitochondrial membrane in certain levels of RSV. This phenotype has also been reported in mouse muscle cells C2C12, when cells were supplemented with 25 μM RSV (Price et al. 2012). In addition, several studies reported that RSV depolarizes the mitochondrial membrane. This effect could be due to a long-term effect of RSV supplementation, occasioned by augmented ROS generation due to an increase of mitochondrial membrane potential and the inhibition of the complex I activity.

Contrary to the traditional perception that RSV is an antioxidant molecule, recently it has been found that RSV actually acts as a pro-oxidant molecule (Plauth et al. 2016). It has been demonstrated that RSV displays a hormetic effect on cell viability dependent on ROS generation (Plauth et al. 2016; Ristow 2014). At high doses of RSV (>50 μM), a drastic decrease in cell viability occurs. Whereas low doses of RSV (<50 μM) induce the proliferation of mammalian cells. Both phenotypes were nullified by GSH addition, a well-known quencher of oxidant molecules (Plauth et al. 2016). Interestingly, we found that RSV supplementation decreases H_2_O_2_ release at high-level glucose, while at low-level glucose increases it. A hormetic pattern in H_2_O_2_ release at low-glucose level with RSV supplementation as it does occur in mammalian cell lines was observed. However, the decrease of H_2_O_2_ release with high-level glucose could be attributed to highly coupled ATP generation by OXPHOS, which corresponds to the increase of oxygen consumption under this growth condition with low electron leakage. Nevertheless, the increase of mitochondrial membrane potential suggests that the electron transport chain could be further reduced, which promotes ROS generation by reverse electron transport. ROS generated might induce antioxidant systems, which in turn reduces H_2_O_2_ release. Consequently, this data indicates that RSV acts as a pro-oxidant molecule with a mitochondrial origin.

In conclusion, this data indicates that RSV decreases the CLS in *S. cerevisiae* by induction of mitochondrial dysfunction in a glucose-dependent manner.

## Acknowledgments

This study was funded by grants from Instituto Tecnológico Superior de Ciudad Hidalgo (3308.100310), Tecnológico Nacional de México (165.14.2-PD and 166.14.2-PD) and Programa para el Desarrollo Profesional Docente (PRODEP program; ITESCH-002). The authors would like to thank M.C Adriana González-Gallardo and Dra. Anaid Antaramian (Unidad de Proteogenómica del INB-UNAM) for their technical support.

